# Decoupling species richness and interaction frequency reveals how fungal interactions regulate wood decomposition

**DOI:** 10.64898/2026.02.18.706504

**Authors:** Yu Fukasawa, Aoi Chiba

## Abstract

Wood decay fungi play a central role in forest carbon cycling, yet the mechanisms linking fungal biodiversity to decomposition remain unclear because species richness and interspecific interactions are rarely separated. Here, these effects were experimentally disentangled using a laboratory wood decomposition microcosm with four common wood-decay fungi. By manipulating the spatial arrangements of pre-colonized wood blocks, fungal species richness and the frequency of interspecific interactions were independently varied. Wood mass loss was quantified, and lignin and carbohydrate analyses were conducted to examine the changes in decay strategies and the potential accumulation of recalcitrant fungal products. Both fungal species richness and interspecific interactions enhanced wood decomposition, but their effects depended on species identity and combinations. Selection effects were observed when competitively dominant species replaced weaker competitors. Several species combinations showed decomposition rates exceeding those of pure cultures, indicating complementarity and facilitation by interspecific interactions. Generalized linear mixed models revealed that interaction frequency and species richness independently influenced decay rates in a species-specific manner. Chemical analyses revealed that interspecific interactions altered the relative loss of lignin and carbohydrates, indicating shifts in the enzymatic allocation and/or production of acid-insoluble fungal metabolites during competition. Our results indicate that competitive interactions among wood decay fungi often accelerate decomposition to offset energetic costs. However, deadlock interactions among basidiomycetes may promote the accumulation of recalcitrant fungal compounds, potentially slowing decomposition over longer timescales. We propose the “accumulated inhibitor hypothesis” to reconcile contrasting fungal diversity–decomposition relationships and highlight the importance of interaction frequency in fungal biodiversity–ecosystem functioning research.

## Introduction

Wood inhabiting fungi play a central role in the global carbon cycle by decomposing dead wood (Bardgett et al. 2008; Bradford et al. 2014), which is a large carbon stock in forest ecosystems (Pan et al. 2011). Decay abilities of wood components, such as lignin and holocellulose, are dependent on fungal species and strains (Schilling et al. 2020). Thus, different fungal communities exhibit distinct wood decay functions (Fukami et al. 2010; Fukasawa and Matsukura 2021). Understanding the relationship between fungal community and wood decay abilities is necessary to predict wood decay functions by fungal communities in changing environment (Lustenhouwer et al. 2020).

Biodiversity–ecosystem functioning (BEF) is an essential topic of ecology in the era of great extinction and climate change (Loreau et al. 2001; Oliver et al. 2015). Several studies on plant communities revealed the positive effects of plant diversity on primary production and associated carbon sequestration (Tilman et al. 2001; Yang et al. 2019; Chen et al. 2025). The BEF theory attributes the effects of diversity to selection and complementarity (Loreau and Hector 2001). The selection effects, which are also referred to as sampling effects, reflect the increased likelihood that diverse communities include highly productive species that disproportionately contribute to productivity (Huston 1997). Compared with selection effects, complementarity effects reflect the positive interactions or niche differentiation among species, which increase productivity beyond what any single species can achieve alone (Cardinale et al. 2007). Complementarity effects may arise not only from resource partitioning but also from facilitative interactions, where one species enhances the performance of others, sometimes referred to as facilitation effects (Callaway and Walker 1997).

BEF debate is also important in the decomposer subsystem (Gessner et al. 2010; Nielsen et al. 2011; Runnel et al. 2025). However, in fungal community, previous BEF studies reported the various effects of fungal species richness on wood decomposition. Several studies reported negative correlations between fungal species richness and wood decay or CO2 emission rates from deadwood in field experiments (Yang et al. 2016; Skelton et al. 2019; Smith and Peay 2021). By contrast, van der Wal et al. (2015) reported a positive relationship between fungal species richness and wood mass loss in field experiments. Other studies reported no correlations, indicating the importance of fungal community identity rather than species richness (Dickie et al. 2012; Yamashita et al. 2015; Venugopal et al. 2017; Jomura et al. 2022). Finding causal relationships between fungal diversity and wood decomposition based on the results of such correlation analyses are always challenging. Smith and Peay (2021) compared the data of field and laboratory incubation experiments and concluded that the results are dependent on the spatial scale: in a micro scale, such as single dead wood, fungal community composition and diversity might drive the decomposition rate, whereas in a larger scale, such as a stand, environmental conditions would be more important. Nevertheless, in microcosm studies with controlled species richness and conditions, the negative (Toljander et al. 2006; Fukami et al. 2010; Fukasawa and Matsukura 2021; Smith and Peay 2021) and positive (O’Leary et al. 2019; Banik et al. 2024) effects of fungal species richness on wood decomposition were reported. Negative relationships seem to have richer evidence so far. However, three of these previous studies referring negative correlations (except for Toljander et al. 2006) evaluated fungal species richness based on the number of fungal species re-detected from the substrates after the incubation experiments. Thus, the researchers cannot reject the possibility that the causality was opposite, that is, slowly decaying substrates host richer fungal communities. Rather, many studies on two-species dual culture reported the positive effects of fungal species interaction on wood decomposition (Hiscox et al. 2015, 2018; Fukasawa and Kaga 2022; Cui et al. 2023). These results indicated that the relationships between fungal species richness and wood decomposition are context dependent.

Daniel Maynard and his colleagues shed light on this complex situation. They coupled incubation experiments of 10–18 wood decay fungi and computer simulation models to show that species diversity alone has negligible impacts on community functioning but the diversity interacts with the key properties of the competitive network, including competitive intransitivity and average competitive ability, to shape biomass production, respiration, and carbon use efficiency and to ultimately affect the direction of diversity–function relationships (Maynard et al. 2017a, b). Interestingly, fungal communities consisting of strong competitors showed negative relationships, whereas those that consist of weak competitors showed positive relationships between species richness and community carbon use efficiency (Maynard et al. 2017b).

Although these two studies were conducted using fungal communities grown on agar media, they extended the system to wood decomposition in the field and found that community evenness is a key mediator between species richness and wood decay rate: highly even communities, consisting of co-existing species, exhibit a positive richness–decay relationship and uneven communities, dominated by strong competitors, exhibiting a negative or null response (Maynard et al. 2018). These results indicated that the quantity of competitive interactions in fungal communities is essential to evaluate the wood decay functions of the communities.

Competitive and antagonistic interactions are energetically expensive for fungi because they produce a variety of enzymes (Baldrian 2004; Gregorio et al. 2006; Hiscox et al. 2010) and secondary metabolites, such as antibiotics (Helaly et al. 2018; Matik et al. 2025), volatiles (O’Leary et al. 2019), and pigments/recalcitrant chemicals (Krause et al. 2020; Dullah et al. 2021; Morris et al. 2021), on competition to inhibit antagonist activities (Hiscox and Boddy 2017). Thus, increased cost due to competition may lead to reduced resource allocation to other activities such as hyphal growth and decomposition by energetic trade-offs (Fukami et al. 2010; Maynard et al. 2017a).

Similarly, frequent competition may lead to the accumulation of antifungal and recalcitrant materials within wood, which could reduce wood decomposition. However, increased competition cost could accelerate resource utilization to compensate for the energy loss (Fukasawa et al. 2020). Therefore, the quantification of competition might be critical for understanding the mechanisms underlying fungal diversity–decomposition relationships. For example, the difference in interaction frequency may affect wood decomposition even in the communities of the same species number of fungi. The separation of selection effects (species number) and complementarity effects (interaction frequency) had long been discussed in general BEF studies (Huston 1997; Loreau and Hector 2001; Chen et al. 2025). However, few studies evaluated the effects of fungal species richness and interaction frequency separately in wood decomposition BEF studies (O’Leary et al. 2021; Banik et al. 2024). This is a bit strange because it is easy to manipulate interaction frequency without changing the numbers of species or genets in fungi as modular organisms (Hiscox et al. 2018). In addition, competition for space among fungal colonies of various sizes is a realistic situation in wood decay fungal communities (Coates and Rayner 1985). Thus, obtaining ecological implications from the results would be valuable.

In the present study, a laboratory decomposition experiment was conducted using the strains of four wood decay fungal species. By varying the combination patterns of wood blocks colonized by these strains, fungal species richness and the frequency of interspecific interactions were independently manipulated. In addition, lignin and carbohydrate analyses were performed on a subset of wood blocks to examine the changes in resource utilization by fungal strains and the potential production of recalcitrant compounds. Based on the results, the effects of fungal species richness and interaction frequency on decomposition, as well as their underlying mechanisms, were disentangled. Although facilitation effects are often included within complementarity effects together with resource partitioning, facilitation effects are frequently treated separately from resource partitioning in decomposition studies (Gessner et al. 2010). This convention was followed in this study.

## Methods

### Fungal strains

Four fungal strains were used in this study (Table S1). All species are common on dead wood of Fagaceae trees, and they cause white rot (or soft rot in the case of Annulohypoxylon truncatum). All strains were maintained on 2% malt extract agar media (MA: 20 g malt extract [Nacalai tesque, Kyoto, Japan], 15 g agar [Nacalai tesque, Kyoto, Japan], 1000 mL distilled water) at 25°C in the dark until use for experiment.

### Competition on agar medium

Prior to the wood block microcosm experiment, the competitive abilities of the four fungal strains were tested on 2% MA arena in Petri dishes with 9 cm diameter. The amount of MA in each dish was 10 mL. The inoculum agar plugs were cut out from the actively growing mycelium on 2% MA using a sterilized cork borer (5 mm diameter). The plugs of different strains were placed on a MA arena at a distance of 3 cm apart from each other. To create a contact of the two strains at the center of the dish, the slow-growing strains were inoculated several days earlier to the fast-growing strains.

The dishes were incubated at 25°C in the dark for 2–4 weeks until one of the two competitors in the dish completely replaced its counterpart (R) or the two competitors in a dish made a zone line at the interaction front and deadlocked (D). The competition of the four strains was settled as a full-factorial design (a total of six combinations), and three replicates were prepared for each combination. To compare the competitive abilities of the four strains, a score of 1 was given to the winner in R dishes, –1 to the loser in R dishes, and 0 to each strain in D dishes. The scores of all dishes were summed up to make the total score of each strain.

### Wood block microcosm

Kiln-dried (70°C) beech (Fagus crenata Blume) wood blocks (1.5 cm ×1.5 cm × 1.5 cm) were numbered by using a B pencil and weighed before use in the experiment. The blocks were soaked in autoclaved deionized water for half day and then autoclaved for 20 min at 121°C. Autoclaving was repeated three times with one day interval to completely sterilize heat-tolerant microbes.

The autoclaved blocks were placed onto the two-week-old mycelium of fungal strains growing on 0.5% MA in PP deli pots (13 × 8 bottom with 5 cm height; KUNGFU Inc., Tokyo, Japan), with 12 blocks per pot. The pots were incubated at 25°C in the dark for two months to allow the strains to colonize the wood blocks. A total of 2240 wood blocks were prepared in 191 pots.

After two-month colonization, the wood blocks were aseptically harvested and used for microcosm experiments. Sixteen colonized wood blocks were arranged 4 × 4 in various combinations of species and patterns in accordance with the experimental design (Fig. 1). We set up experiments including single species (pure culture), two species, and four species. In pure culture, four experiments were set up because we had four fungal species. In two-species experiments, three patterns of the 16-block arrangement were prepared to make 4, 8, and 24 interaction surfaces. An interaction surface (shown as white dots in Fig. 1) consisted of two wood blocks with different fungal species facing each other. Each pattern had six experiments because there are six ways to choose two species from four. In four-species experiments, arrangements with only 8 and 24 interaction surfaces were prepared because of mathematical limitation.

**Fig. 1.**
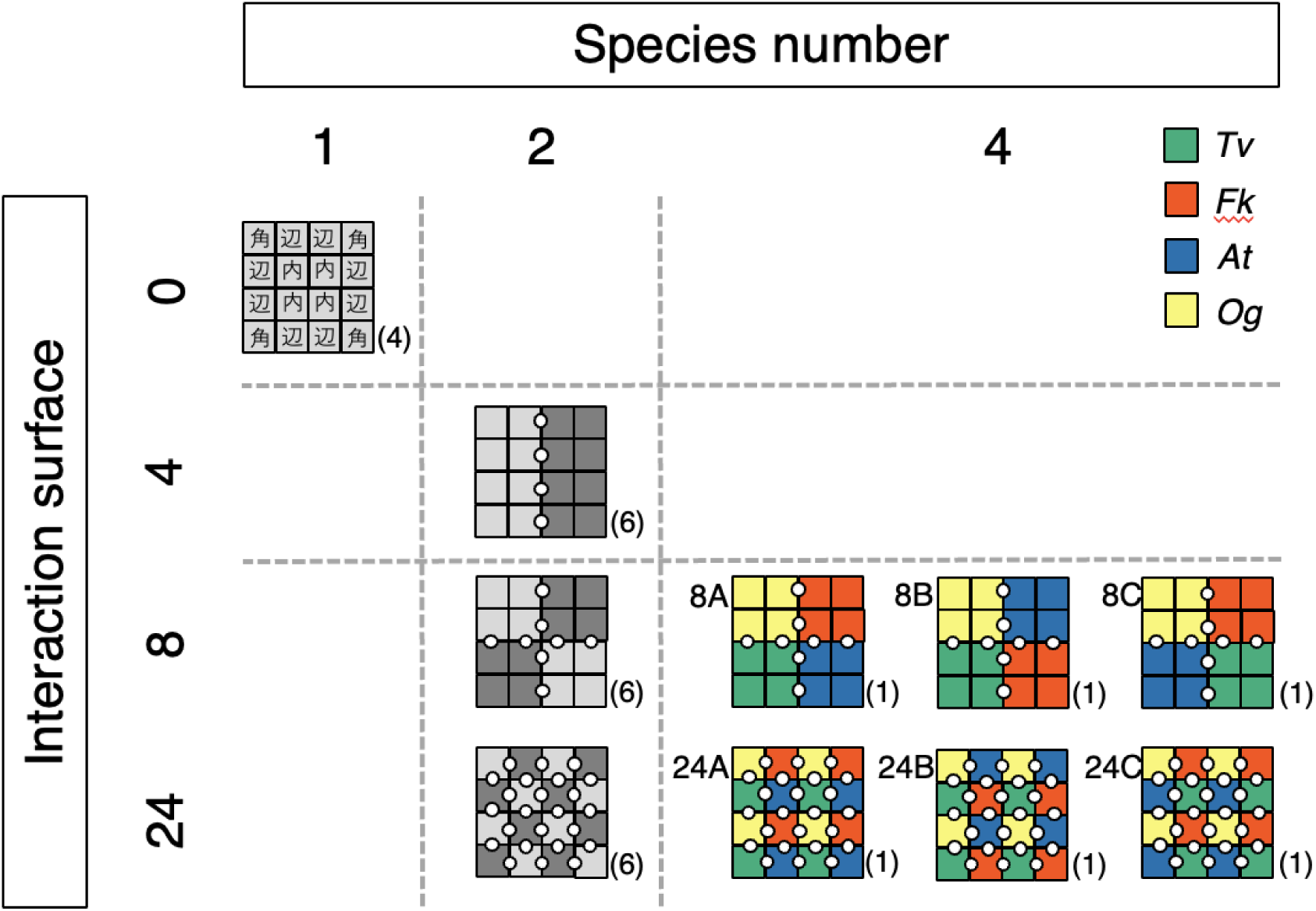
Experimental design of wood block microcosms. The number of fungal species was 1 (pure culture), 2, and 4. In mixed cultures, the interaction frequency (number of interaction surface) was 4 (only in two-species combinations), 8, and 24. An interaction surface (shown as white dots in the figure) consists of two wood blocks with different fungal species facing each other. In a four-species combination, three different arrangements of species location were prepared for each of the 8 and 24 interaction frequencies. The letters assigned to the blocks in pure culture indicate the location in the experiment: 角, corner; 辺, side; 内, center. A total of 28 experiments were designed, and five replicate pots were prepared for each experiment.

Instead, three options that differ in fungal species location were prepared for each of the 8 and 24 interaction surface arrangements. In total, 28 experiments differ in the number of species, interaction surface, and species combinations and locations were prepared.

Each experiment had five replicates. In each replicate, the 16 wood blocks were bound using a sterilized PE strand to keep in contact with one another and were placed on 150 mL of autoclaved perlite in a PP deli pot (13 cm × 8 cm × 5 cm). All wood blocks were positioned to make their vessels vertically directed because the direction of wood vessels can affect the outcome of fungal competition (O’Leary et al. 2018). In addition, 25 mL of autoclaved deionized water was added to perlite for moisture. Five small holes (1 mm diameter) were made on the pots covered by a microporous surgical tape (COME’S, Hyogo, Japan) to allow aeration. A total of 140 pots were covered with a lid, incubated at 25°C in the dark for another three months, and watered once at 1.5 months after the start of incubation to maintain moisture.

At the end of the experiment, the blocks were separated, and fungal hyphae and perlites attached to the surface were removed using a razor blade. Then, the blocks were weighed fresh and vertically split in half using a sterile chisel in laminar flow hood. A half of the wood block was weighed fresh again and then kiln dried at 70°C for over two weeks until a constant weight is obtained. Then, these weights were used to calculate the water content of each block and to estimate the dried weight of each wood block after the incubation period using the following equations:

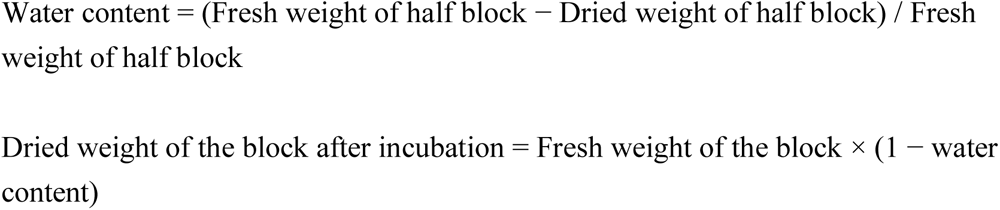

Furthermore, the weight loss (%) of each block after the incubation experiment was calculated as the percentage of weight loss against the original dry weight of the block.

From another half of the block, a piece of wood (approximately 5 mm^3^) was excised from the center of the split surface, placed onto 2% MA and incubated at 25°C until the mycelium had emerged and could be identified morphologically. The competition outcomes were recorded as whether the fungal species replacement from the original wood colonizers occurred or not. Furthermore, the presence of zone lines was recorded if they appeared.

### Chemical analysis of wood blocks

The lignin and carbohydrate contents of the wood blocks originally colonized by *Fuscoporia koreana* (hereafter referred to as *Fk*) or *Omphalotus guepiniformis* (hereafter referred to as *Og*) were further analyzed after the incubation experiment if they were not replaced with other species nor had zone lines. Among the *Fk* wood blocks that meet the abovementioned criteria, 48 blocks in randomly selected three packs of five pure culture packs, 72 blocks incubated with *Og* blocks, 72 blocks incubated with *Trametes versicolor* (hereafter referred to as *Tv*) blocks, 72 blocks incubated with *A. truncatum* (hereafter referred to as *At*) blocks, and 72 blocks in four-species packs were used for chemical analysis (a total of 336 blocks). Among the *Og* wood blocks that meet the abovementioned criteria, 48 blocks in randomly selected three packs of five pure culture packs, 13 blocks incubated with *Fk* blocks, 109 blocks incubated with *At* blocks, and 91 blocks incubated with *Tv* blocks were used for chemical analysis (a total of 261 blocks). The *Og* blocks in four-species packs were not used for chemical analysis because of their limited number.

The dried split half of these blocks were further cut into finer particles and then pulverized to pass through a 0.5-mm screen using Multi-beads Shocker (MB3200 (S), Yasui Kikai, Osaka, Japan) for chemical analysis. The amount of Klason lignin in the sample was estimated gravimetrically using hot sulfuric acid digestion (King and Heath 1967). The samples were extracted with alcohol-benzene at room temperature, and the residues were treated with 72% sulfuric acid (v v^−1^) for 2 h at room temperature with occasional stirring. Then, the mixture was diluted with distilled water to make a 2.5% sulfuric acid solution and autoclaved at 120°C for 60 min. After cooling, the residue was filtered and washed with deionized water through a porous crucible (G4), dried at 105°C, and weighed. The filtrate (autoclaved sulfuric acid solution) was used for total carbohydrate analysis by using the phenol–sulfuric acid method (Dubois et al. 1956).

Five percent phenol (v v^−1^) and 98% sulfuric acid (v v^−1^) were added to the solution. The optical density of the solution was then measured by using a spectrophotometer at 490 nm using the known concentrations of D-glucose as standards. Considering that wood contains very little carbohydrate other than holocellulose, the measured amounts of total carbohydrate were regarded as holocellulose contents in this study.

After acid hydrolysis, the remaining residue contains a mixture of organic compounds including not only lignin, but also condensed tannins and waxes (Preston et al. 1997). In addition, Fukasawa et al. (2009) reported that the net amount of Klason lignin fraction had increased after the competition between fungal colonizers on twig litter. These results indicated that the Klason lignin measured in this study includes acid-insoluble components produced by fungi.

Lignin loss (%)/wood loss (%) rate (L/W) and carbohydrate loss (%)/wood loss (%) rate (C/W) are useful indices of the relative decay rate of wood fractions (Fukasawa et al. 2009), and were calculated.

### Data analysis

All statistical analyses were performed using R version 4.3.1 (R core team 2023).

The weight loss and water content of wood blocks were compared across fungal species, block positions, and experiments using the Steel–Dwass test (*NSM3* package, *pSDCFlig* command, Asymptotic) (Schneider et al. 2023). A Chi-square test was used to compare the reisolation frequency of fungal species across the experiments. Adjustment of *p* values in multiple comparisons was performed by using the Benjamini–Hochberg method.

To determine the factors that are associated with the weight loss of the wood blocks of each fungal species, generalized linear mixed models (GLMMs) were performed using the lme4 (Bates et al. 2022) and lmerTest packages (Kuznetsova et al. 2016). For this analysis, the data of the wood blocks in which the original colonizer fungus was reisolated without replacement by other fungal species were used to confirm that the wood decay was certainly due to that fungus. Considering that all the *At* blocks were replaced by other fungal species, the data of *At* blocks were not included in this analysis. In the models, the dried weight of the wood blocks after the incubation experiment was set as a responsive variable, pack ID as a random effect, and the original dried weight of the wood blocks as an offset term. The explanatory variables included the number of surfaces faced to the block of other species, the number of surfaces faced to the block of the same species, the number of fungal species included in the pack, and the water content of the wood blocks. The number of surfaces faced to the block of the same species was included in the model to distinguish the situation from the block surfaces faced to the air. Model selection was performed using the *dredge* command in the *MuMIn* package (Burnham and Anderson 2002; Bartoń 2025) based on the Akaike information criterion. For the number of fungal species in the pack, a model that includes either of the number before the incubation or after the incubation was made to check the difference in results. The multicollinearity of the variables was checked by using the *vif* command in the *car* package (Fox and Weisberg 2018).

## Results

### Competitive abilities on agar media

The competition outcomes of the four fungal species on 2% MA are shown in Table S2 and Fig. S1. *Og*, *Tv*, and *Fk* were deadlocked to one another. *Fk* was also deadlocked with *At*. *Og* and *Tv* replaced *At*. All three replicate dishes showed the same outcome in all competition. Based on these results, the competition scores were 3 for *Og* and *Tv,* 0 for *Fk*, and −6 for *At*.

### Competition outcomes on wood blocks

Fig. 2 shows the fungal species reisolated from the wood blocks after the incubation experiments. Among the two-species systems (Fig. 2A), *Fk* replaced all the blocks of *Og* when the interaction surface was 4 or 8, but *Og* remained in some blocks when the interaction surface was 24. All the *At* blocks were replaced by all the competitors. *Tv* versus *Og* and *Tv* versus *Fk* were deadlocked, and little change was observed in fungal occupancy in the blocks after the incubation experiment.

**Fig. 2.**
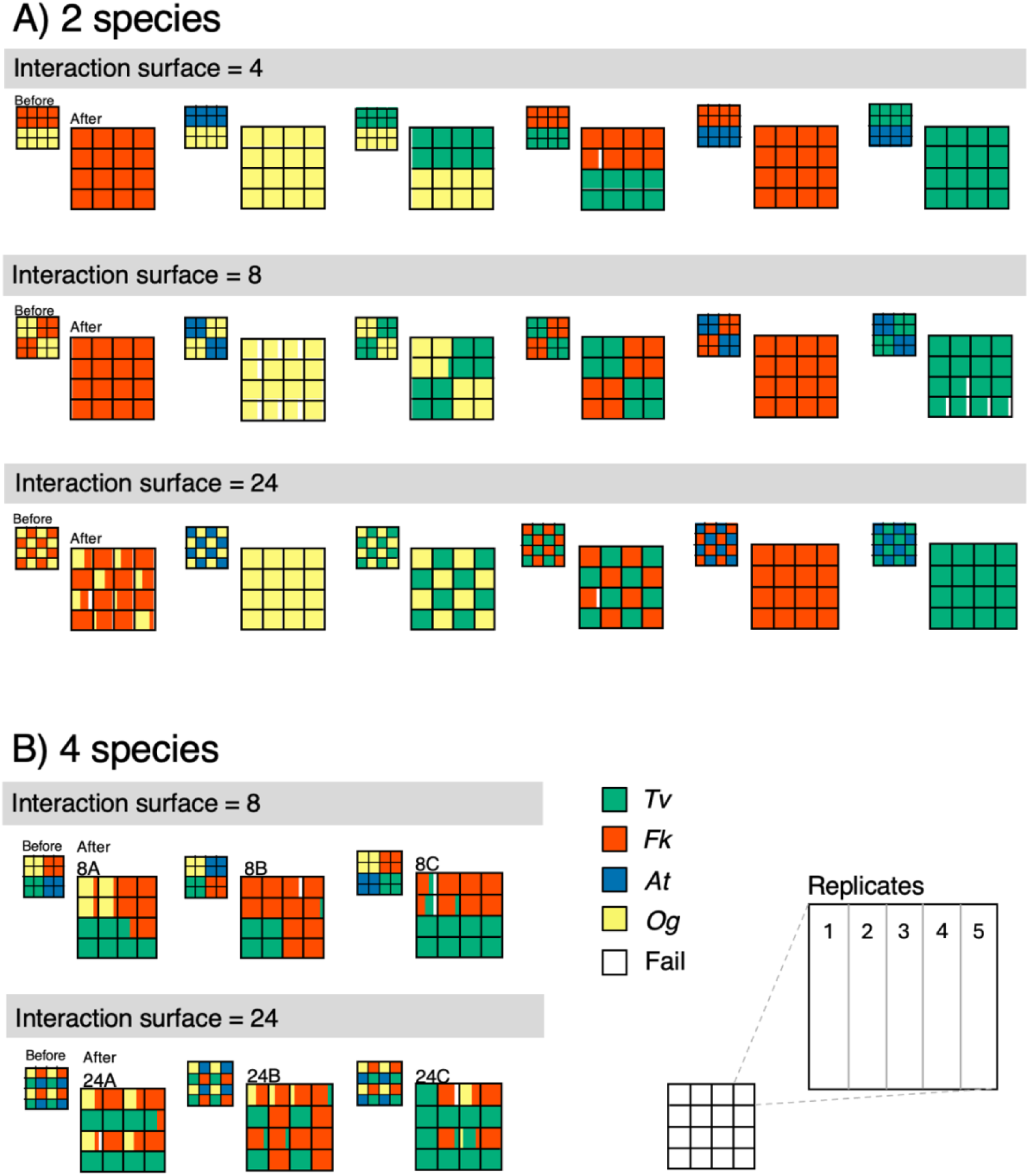
Fungal species reisolated from wood blocks after the incubation experiment of two-species (A) and four-species (B) combinations. The small figure on the upper left of each figure shows the original distribution of fungal species before incubation. Each compartment in the figure is further divided into five segments representing the five replicates, as shown in the bottom right.

The outcome of four-species competition differed across the options of different fungal species locations in 8 and 24 interaction surfaces (Fig. 2B). In 8A and 24A experiments, *Fk* and *Tv* largely invaded into *At* blocks, but the invasion to *Og* blocks was small. In 8B and 24B, only *Fk* invaded into *Og* blocks through *At* blocks. However, in 8C and 24C, only *Tv* invaded into *Og* blocks through *At* blocks. Notably, none of the four-species experiments retained their original four-species sets after the incubation because of the competitive exclusion of *At* from the system.

Table S3 shows that *Fk* occupied the largest number of blocks in most of the experiments, followed by *Tv* and *Og*. *At* lost all of its original wood blocks. In the four-species experiments with 24 interaction surfaces, *Tv* occupied wood blocks more than *Fk*. The Chi-square test indicated that *Tv* showed significantly larger (*p* < 0.05) occupancy in four-species experiments with 24 interaction surfaces than expected. *Fk* showed significantly larger occupancy in four-species experiments with eight interaction surfaces and smaller occupancy in two-species experiments with 24 interaction surfaces than expected. *Og* showed significantly larger occupancy in two-species experiments with four and 24 interaction surfaces and smaller occupancy in four-species experiments with eight and 24 interaction surfaces than expected.

### Wood decay abilities of the four fungal species in pure culture

The weight loss (%) of the wood blocks in pure cultures was significantly different across the four species (Fig. 3A). *Tv* showed the largest weight loss, whereas *At* showed the smallest. In *At*, *Og*, and *Tv*, blocks of inner position (内) showed a larger weight loss than blocks of side (辺) and corner (角) positions. In *Fk*, the weight loss of the blocks was not different across the block positions.

**Fig. 3.**
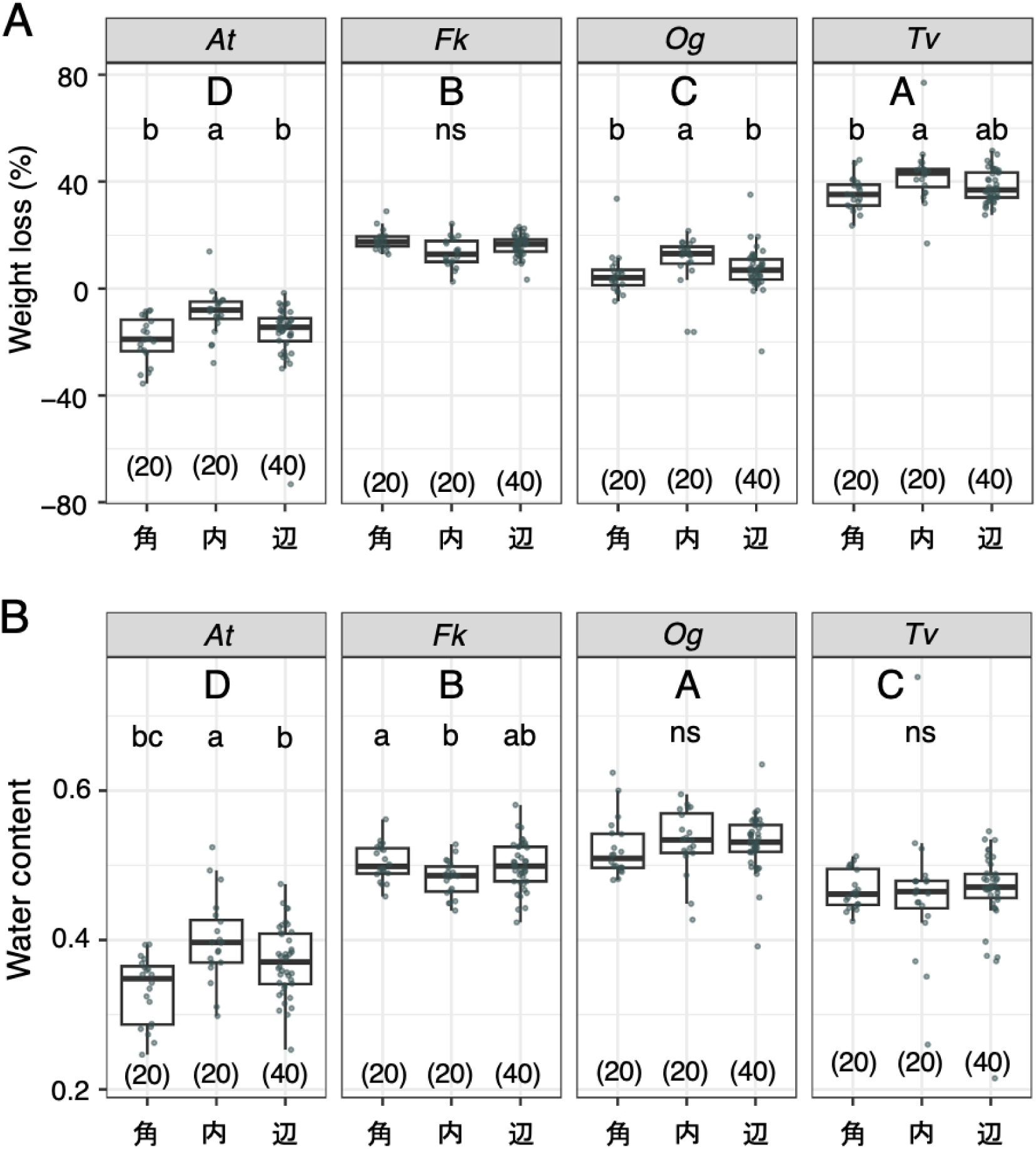
Weight loss (A) and water content (B) of the wood blocks in pure cultures of the four fungal species. The *x*-axis represents the wood block positions shown in Fig. 2: 角, corner (*n* = 20); 内, inner (*n* = 20); 辺, side (*n* = 40). The different uppercase and lowercase letters indicate significant difference across the four species and across the three positions within a species, respectively (Steel–Dwass test, *p* < 0.05). ns, not significantly different. The numbers enclosed in parentheses are the sample number.

The water content of the wood blocks in pure cultures was significantly different across the four species (Fig. 3B). *Og* showed the largest water content, whereas *At* showed the smallest. In *At*, blocks of inner position (内) showed larger water content than blocks of side (辺) and corner (角) positions. In *Fk*, blocks of corner position (角) showed larger water content than blocks of inner position (内). No difference in water content was found across block positions in *Tv* and *Og*.

### Wood decay in species mixture experiments

The weight loss of the wood blocks increased with fungal species number before incubation (Fig. 4A). If the species number after incubation was considered, then weight loss was the largest in two species than one and three species (Fig. S2). Notably, none of the four-species experiments retained their original four-species sets after the incubation. In addition, interspecific interaction positively affected weight loss, whereas the difference in interaction surface had little effect (Fig. 4B). Theh weight loss of the wood blocks in two-species experiments is shown in Fig. 5. In *Fk* versus *Og*, the blocks in the experiment with eight interaction surfaces showed a significantly larger weight loss than those in pure cultures of both competitors. Among the three interaction surfaces, the 8 and 24 interaction surfaces showed a larger weight loss than the four interaction surface. In *At* versus *Og*, all mixed cultures showed a larger weight loss than pure cultures of both competitors, whereas no difference was found among the three interaction surfaces. In *At* versus *Fk*, the weight loss of the wood blocks in mixed cultures did not differ from that of *Fk* pure culture, and no difference was found among the three interaction surfaces. In other combinations, the weight losses of the wood blocks in mixed cultures were intermediate values between those of the two competitors and were not different among the three interaction surfaces. The weight losses of the wood blocks in four-species experiments showed the intermediate values between *Tv* and *Fk*, and such weight losses did not differ among the six experiments with different interaction surfaces and species location (Fig. 6).

**Fig. 4.**
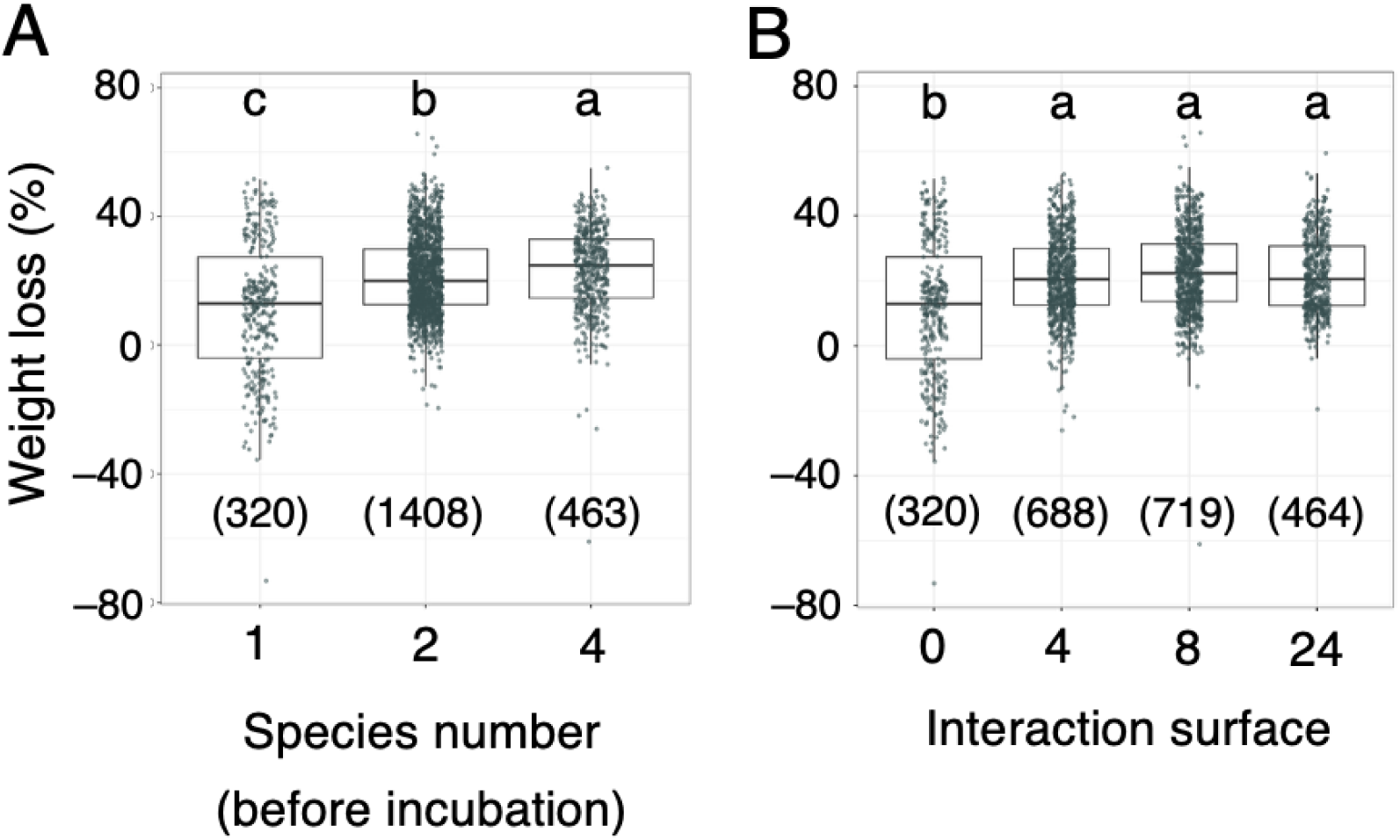
Relationships between weight loss (%) and species number before incubation (A) and interaction surface (B) of the experiments. The different uppercase and lowercase letters indicate significant difference across the three levels of species number (A) and across the four levels of interaction surface (B), respectively (Steel–Dwass test, *p* < 0.05). The numbers enclosed in parentheses are the sample number.

**Fig. 5.**
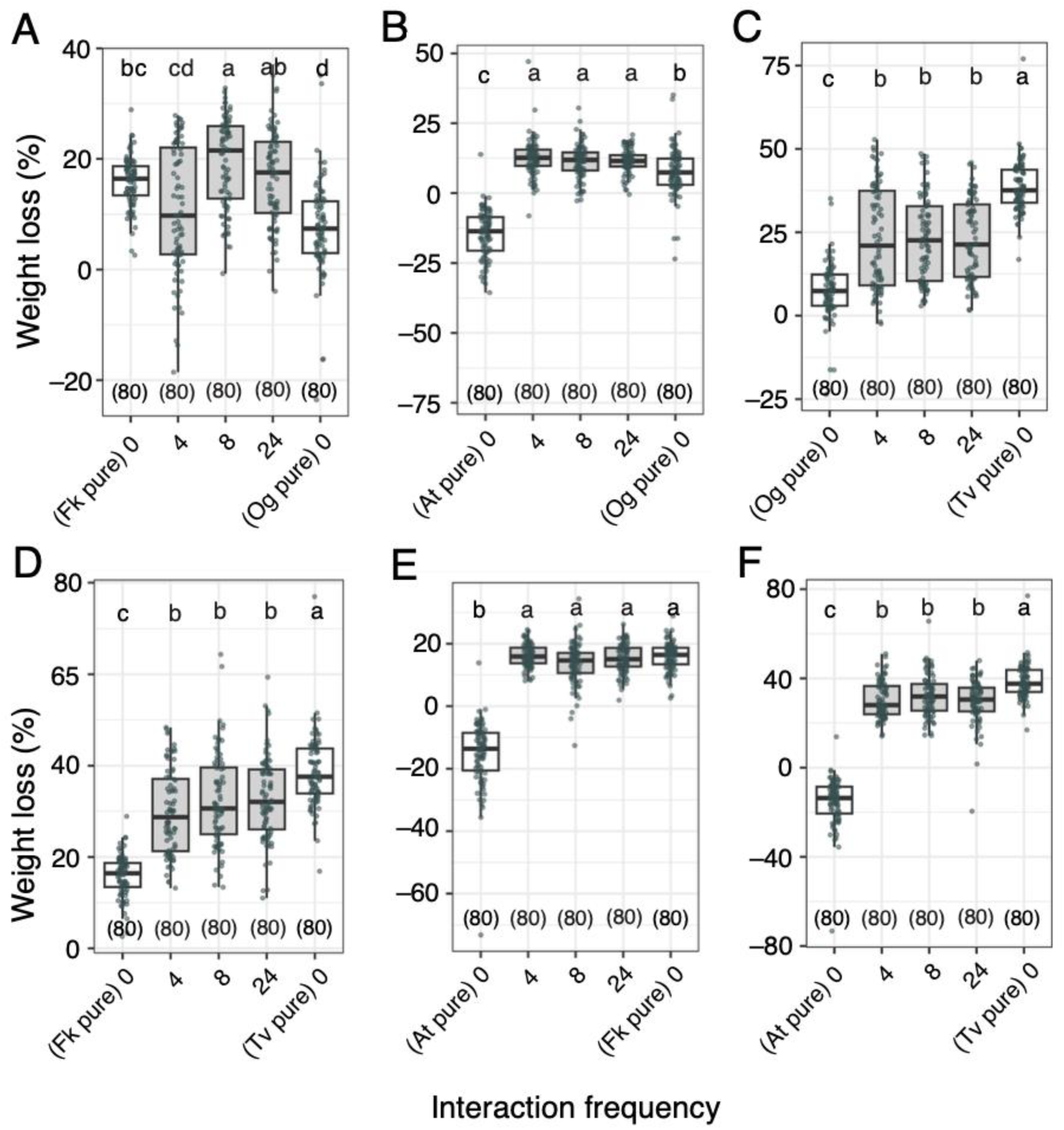
Weight loss of the wood blocks in two-species experiments. (A) *F. koreana* versus *O. guepiniformis*. (B) *A. truncatum* versus *O. guepiniformis*. (C) *O. guepiniformis* versus *T. versicolor*. (D) *F. koreana* versus *T. versicolor*. (E) *A. truncatum* versus *F. koreana* . (F) *A. truncatum* versus *T. versicolor*. In each figure, the white boxes at the left and right sides show the results of pure cultures of the focal species for comparison purposes, and the central three gray boxes show the results of mixed cultures representing the difference in the three interaction frequencies (4, 8, and 24). Blocks of different positions are pooled. The different letters shown above each box indicate significant difference within each figure (Steel–Dwass test, *p* < 0.05). The numbers enclosed in parentheses are the sample number.

**Fig. 6.**
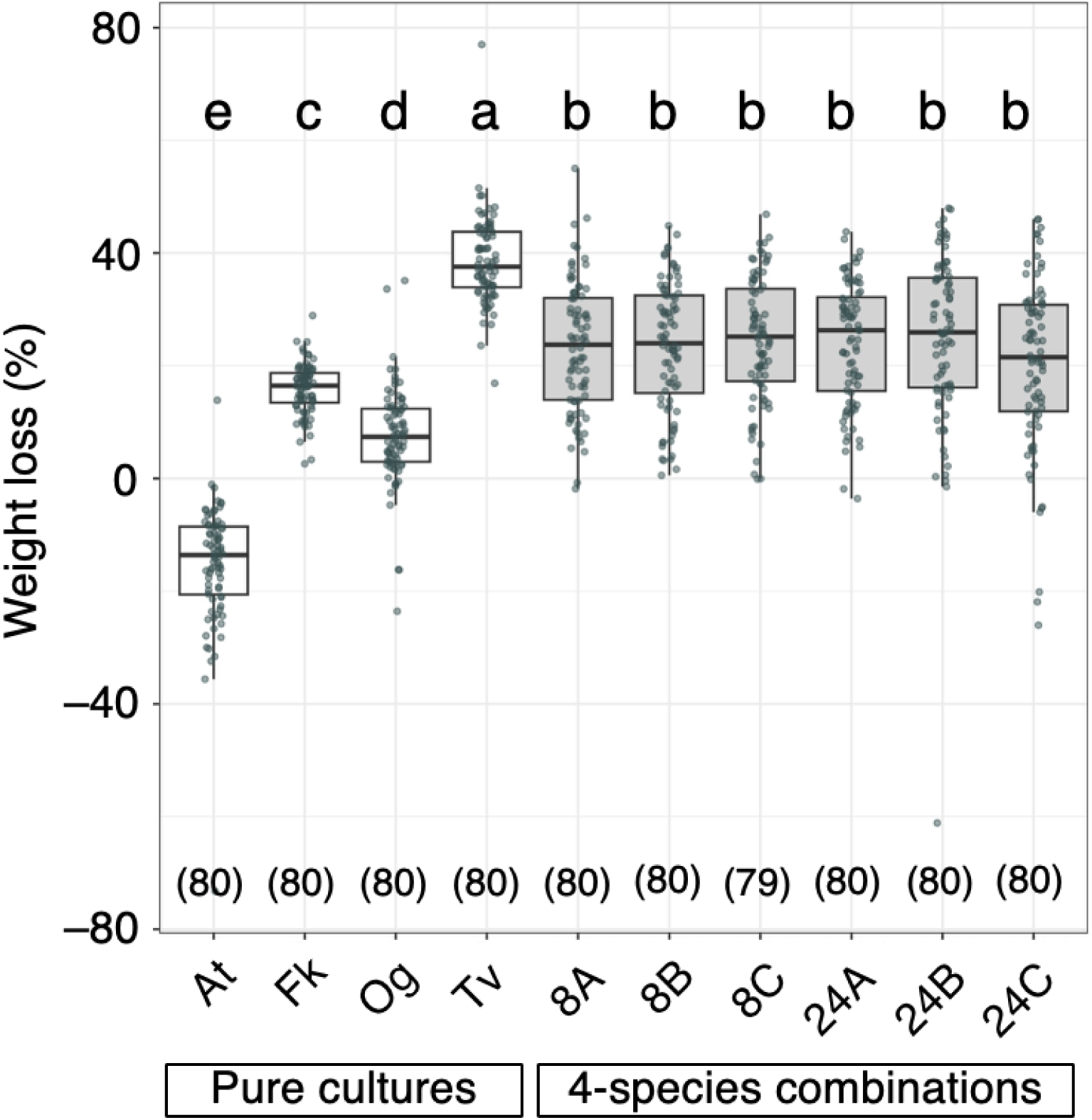
Weight loss of the wood blocks in four-species experiments. The four white boxes at the left show the results of pure cultures, and the six gray boxes at the right show the results of mixed cultures. See Fig. 2 for the species locations in the mixed cultures. The different letters shown above each box indicate significant difference (Steel–Dwass test, *p* < 0.05). The numbers enclosed in parentheses are the sample number.

Among the wood blocks where originally colonized fungus was replaced by the competitor, *At* blocks replaced by *Og* showed a significantly larger weight loss than pure cultures of both fungi in two- and four-species experiments (Fig. 7). Similarly, *At* blocks replaced by *Fk* showed a significantly larger weight loss than pure cultures of both fungi in four-species experiment. In other combinations of species’ replacement, wood blocks showed an intermediate weight loss between the weight losses of the pure cultures of both competitors.

**Fig. 7.**
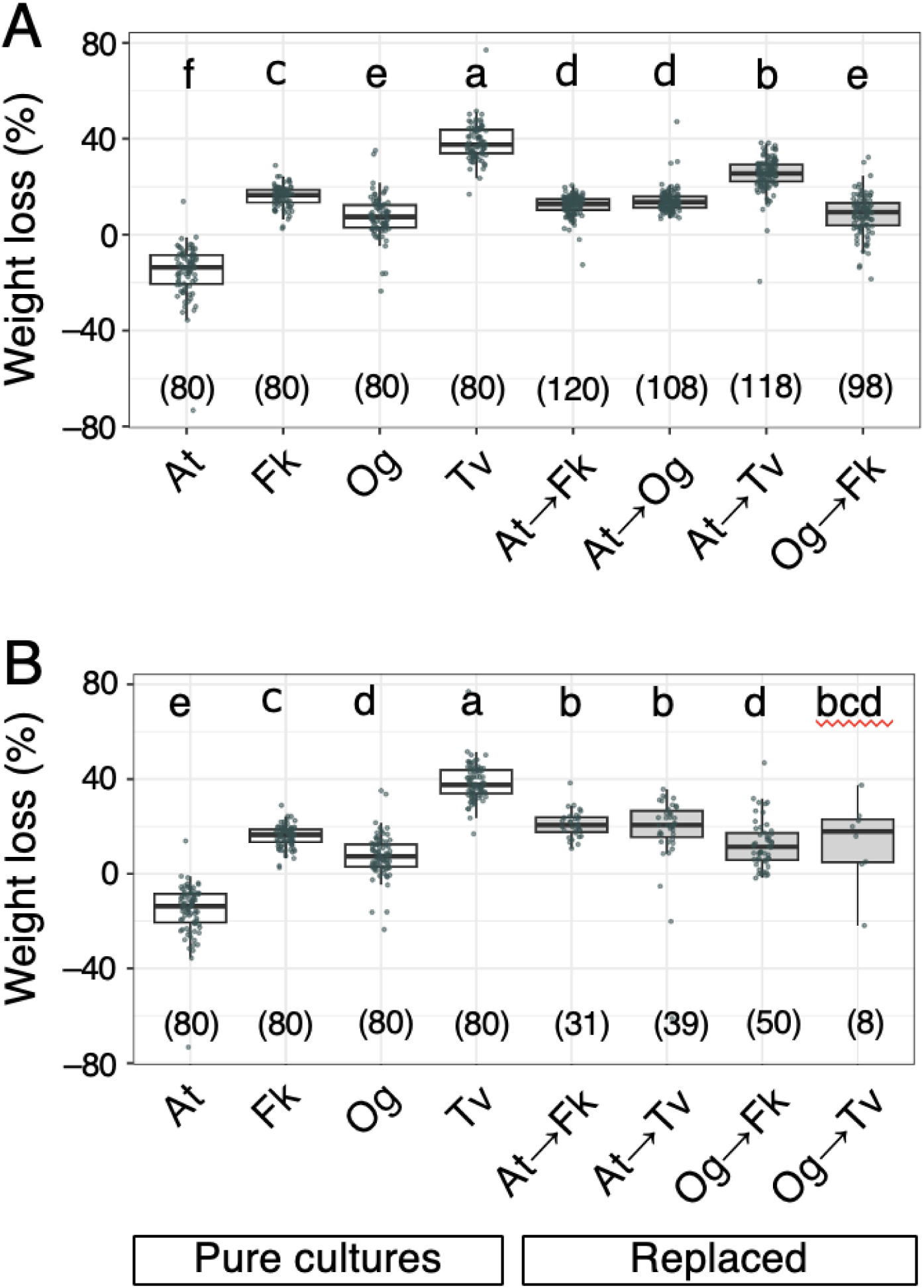
Weight loss of the wood blocks where the species replacement was observed (gray boxes in the right). (A) Two-species experiments. (B) Four-species experiment. The four white boxes at the left show the results of pure cultures for comparison purposes. The different letters shown above each box indicate significant difference (Steel–Dwass test, *p* < 0.05). The wood block of *F. koreana* replaced by *T. versicolor* was only one; thus it was not included in the Steel–Dwass test. The numbers enclosed in parentheses are the sample number.

### Factors associated with weight loss of non-replaced blocks

GLMM revealed that the dry weight of *Tv* wood blocks after the incubation experiment was negatively associated with the number of surfaces that contacted with the same species (i.e., *Tv*) blocks (Table S4). The weight of *Og* blocks was negatively associated with the water content of the blocks and the number of surfaces contacted with other species blocks. The weight of *Fk* blocks was negatively associated with the water content of the blocks and the number of fungal species in the system before incubation. Models used the number of fungal species after the incubation gave similar results (Table S5).

### Lignin and carbohydrate loss in *Fk* blocks

In two-species experiments, the weight loss of lignin and carbohydrate was affected by the competitor species (Fig. 8AB). In *Fk* blocks connected to *At* blocks, the weight loss of lignin and carbohydrate was not different from that of pure culture (Fig. 8A).

**Fig. 8.**
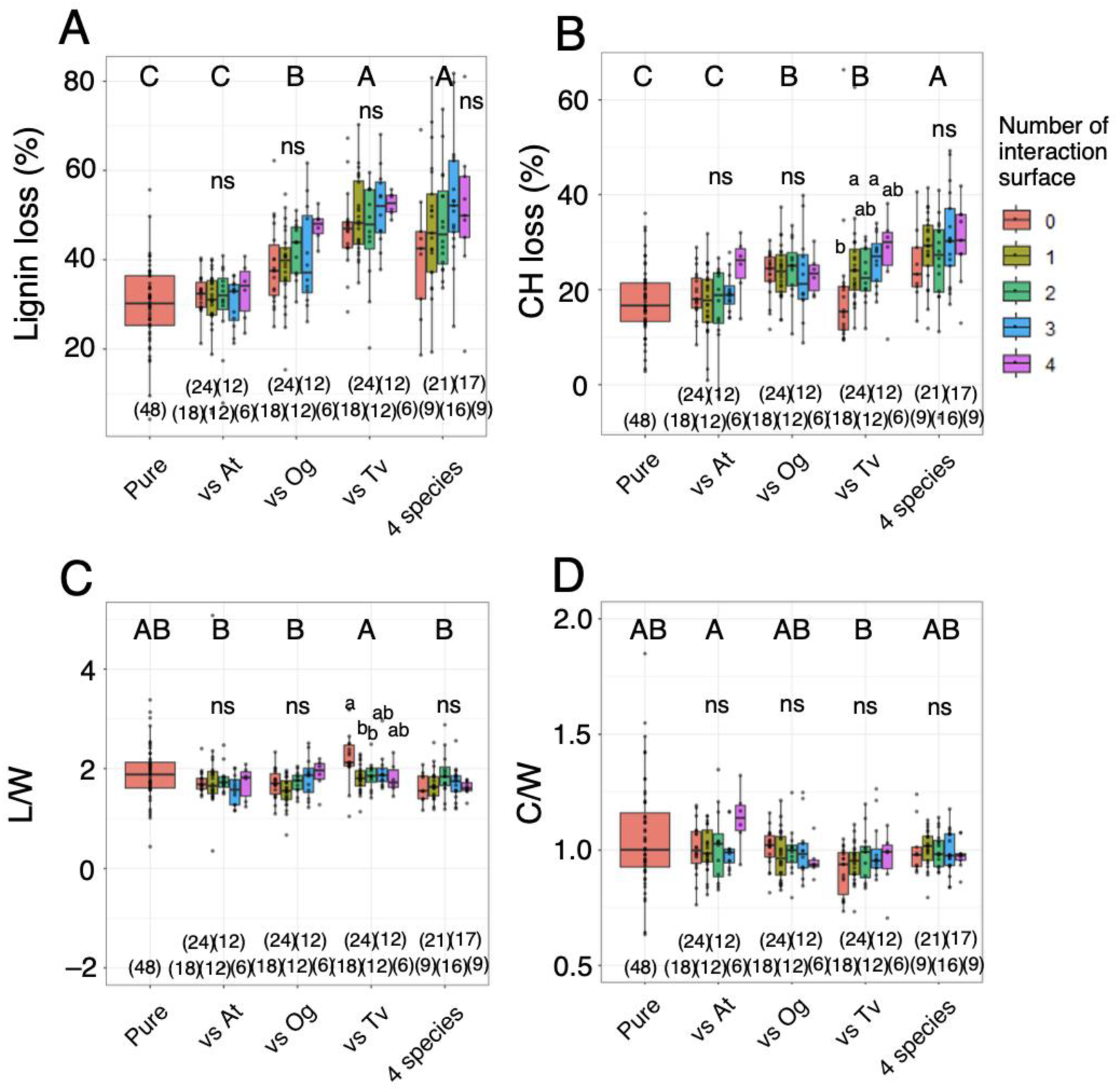
Weight loss (%) of lignin (A) and carbohydrate (B) in *Fk* blocks. Relative weight loss of lignin (C) and carbohydrate (D) against weight loss of wood. The data of 336 *Fk* blocks without replacement and zone lines were used. Two outliers in each L/W and C/W were out of the plots. The different uppercase letters indicate significant difference across the experiments (Steel–Dwass test, *p* < 0.05). The different lowercase letters indicate significant difference across the blocks with different number of surface contact to different species within each experiment (Steel–Dwass test, *p* < 0.05). ns, not significantly different. The numbers enclosed in parentheses are the sample number.

However, *Fk* blocks connected to *Og* and *Tv* blocks showed a significantly higher weight loss of lignin and carbohydrate than the pure culture. The weight loss of lignin in *Fk* blocks connected to *Tv* was the highest, which was not significantly different from that of blocks in four-species experiment. The weight loss of carbohydrate was the largest in the blocks in four-species experiment. The number of surfaces connected to different species also affected weight loss but only in carbohydrate in blocks incubated with *Tv* blocks (Fig. 8B). Blocks with one and three surfaces connected to *Tv* blocks showed a significantly higher weight loss of carbohydrate compared with blocks without connection to *Tv* blocks. The weight loss of lignin did not differ across the number of surfaces connected to different species in any experiments.

L/W and C/W of *Fk* blocks were also affected by the competitor species (Fig. 8CD).

*Fk* blocks connected to *Tv* blocks showed higher L/W and lower C/W than those connected to *At* and *Og* and those in four-species experiment. Among the *Fk* blocks in two-species experiment with *Tv*, the blocks without contact to *Tv* showed significantly higher L/W than those with contact to *Tv* (Fig. 8C). C/W did not differ across the number of surface contact to different species in any experiments.

Lignin and carbohydrate weight losses increased with species number in a pack before incubation, whereas L/W decreased with species number (Fig. 9). C/W was not affected by species number in a pack. However, based on the species number in a pack after incubation, L/W was not responding to species number, and C/W was significantly lower in packs with two species than in packs with one species (Fig. S3).

**Fig. 9.**
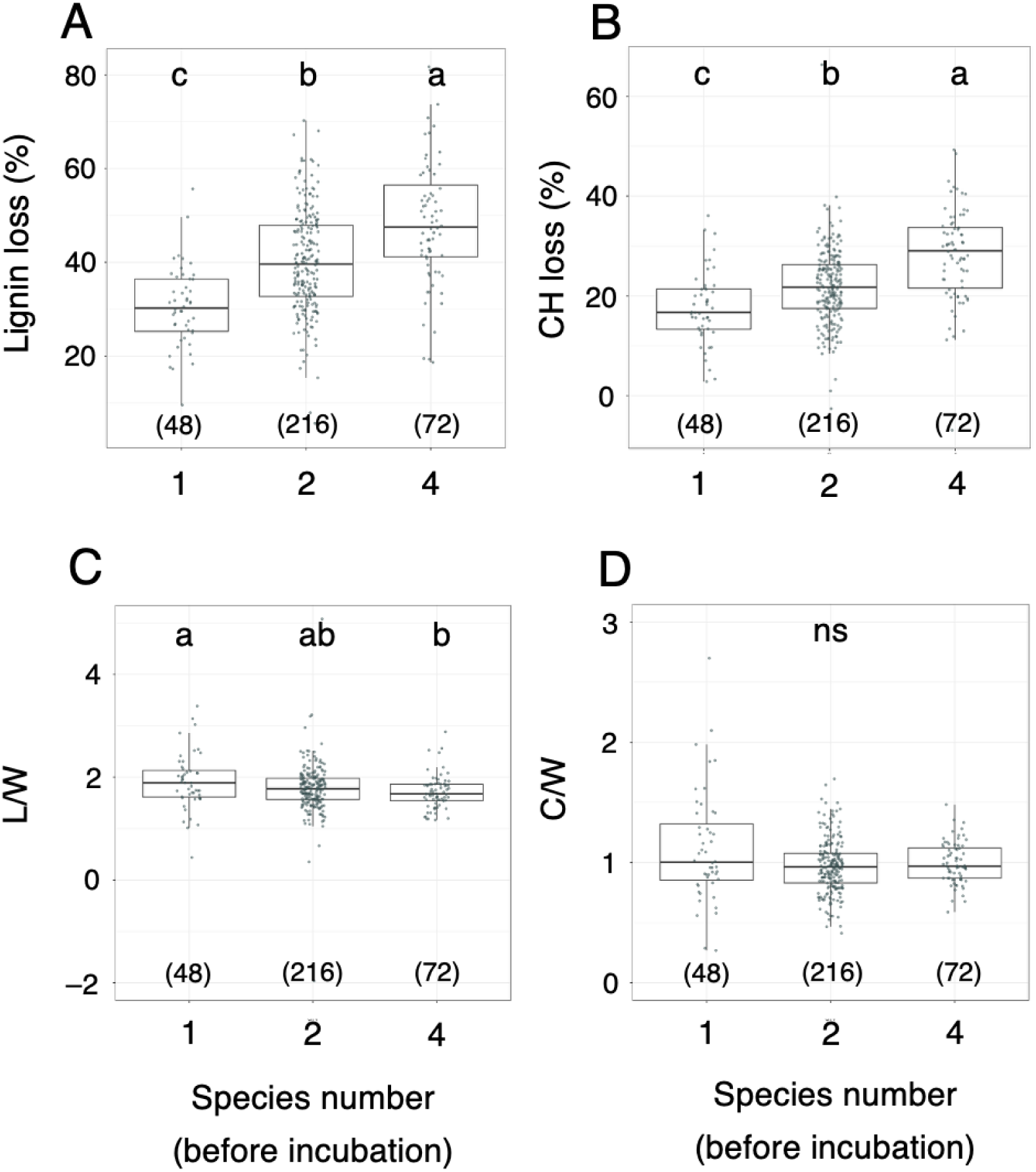
Effects of fungal species number before incubation on the weight loss (%) of lignin (A) and carbohydrate (B) in *Fk* blocks, and the relative weight loss of lignin (C) and carbohydrate (D) against weight loss of wood. The data of 336 *Fk* blocks without replacement and zone lines were used. Two outliers in each L/W and C/W were out of the plots. The different lowercase letters indicate significant difference across the blocks with different fungal species number (Steel–Dwass test, *p* < 0.05). ns, not significantly different. The numbers enclosed in parentheses are the sample number.

### Lignin and carbohydrate loss in *Og* blocks

In four-species experiments, the numbers of *Og* blocks without replacement and zone lines was very limited; thus, they were not used for chemical analysis. In two-species experiments, *Og* blocks connected to *Tv* blocks showed a significantly higher weight loss of lignin than pure culture and blocks incubated with *Fk* (Fig. 10A). The weight loss of carbohydrate, L/W, and C/W were not significantly different across the experiments with different competitor species.

**Fig. 10.**
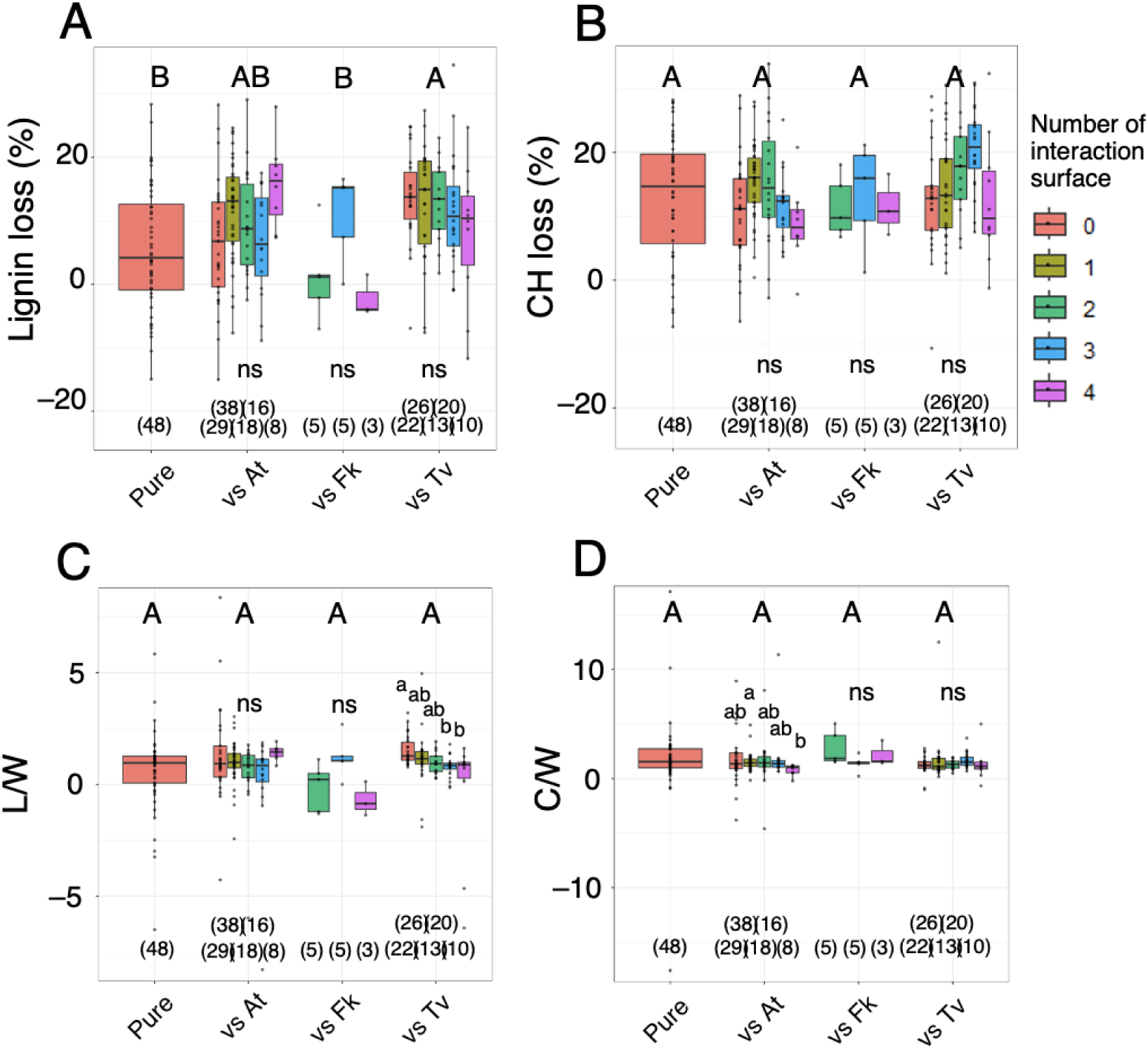
The weight loss (%) of lignin (A) and carbohydrate (B) in *Og* blocks. Relative weight loss of lignin (C) and carbohydrate (D) against weight loss of wood. The different uppercase letters indicate significant difference across the experiments (Steel–Dwass test, *p* < 0.05). The data of 261 *Og* blocks without replacement and zone lines were used. Four outliers in each L/W and C/W were out of the plots. The different lowercase letters indicate significant difference across the blocks with different number of surface contact to different species within each experiment (Steel–Dwass test, *p* < 0.05). ns, not significantly different. The numbers enclosed in parentheses are the sample number.

The number of surface contact to different species had no effects on the weight losses of lignin and carbohydrate (Fig. 10AB). However, L/W of *Og* blocks decreased with the number of interaction surface when incubated with *Tv* blocks (Fig. 10C).

Similarly, C/W of *Og* blocks decreased with the number of interaction surface when incubated with *At* blocks (Fig. 10D).

Only lignin weight loss increased with species number in a pack before incubation, whereas carbohydrate loss, L/W, and C/W did not change between species numbers 1 and 2 (Fig. 11). Based on the species number in a pack after incubation, the results were the same (Fig. S4).

**Fig. 11.**
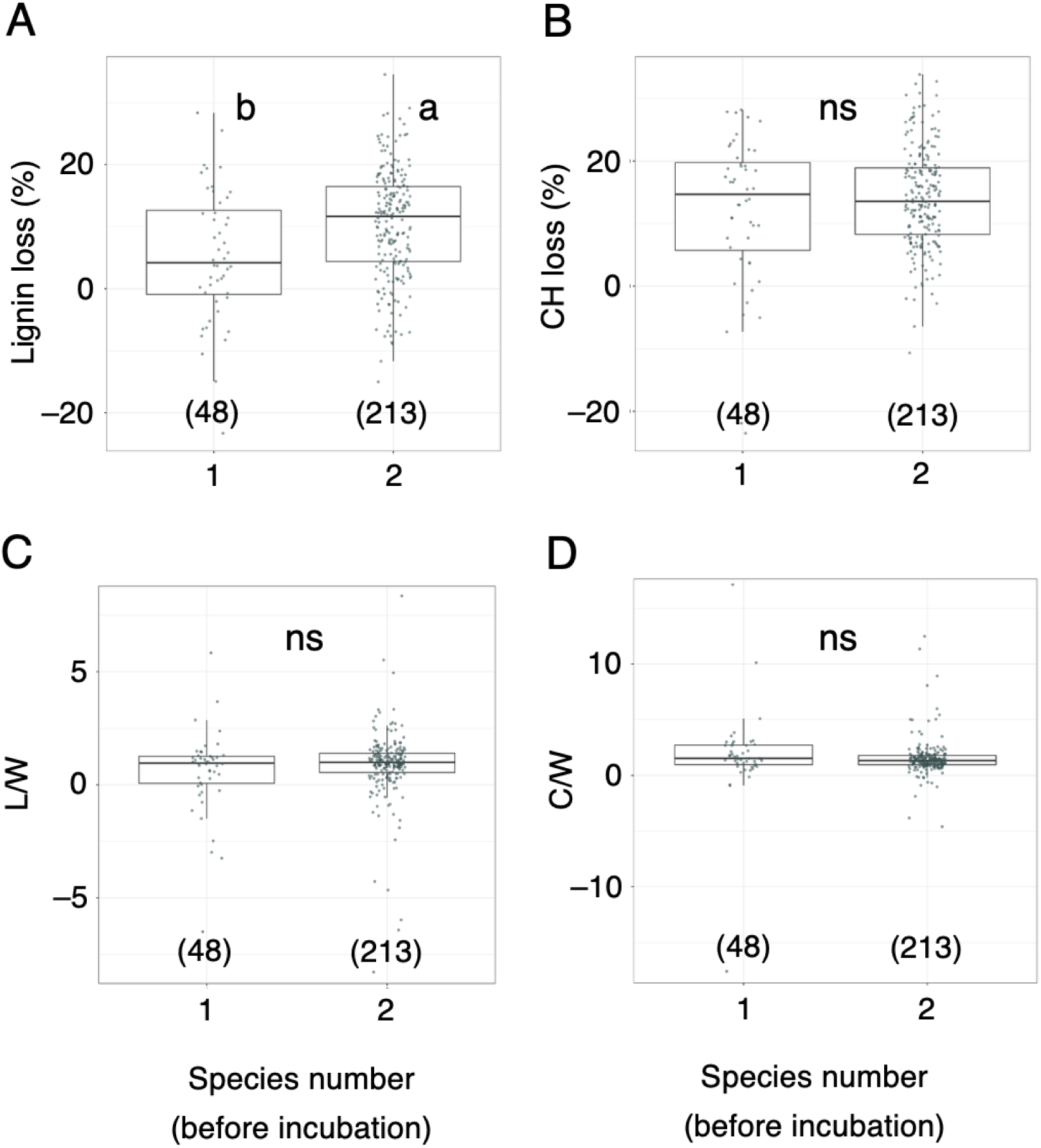
Effects of fungal species number before incubation on the weight loss (%) of lignin (A) and carbohydrate (B) in *Og* blocks, and the relative weight loss of lignin (C) and carbohydrate (D) against weight loss of wood. The data of 261 *Og* blocks without replacement and zone lines were used. Four outliers in each L/W and C/W were out of the plots. The different lowercase letters indicate significant difference across the blocks with different fungal species number (Steel–Dwass test, *p* < 0.05). The numbers enclosed in parentheses are the sample number. ns, no significant difference.

## Discussion

The present study clearly showed that species number and interspecific interaction promoted wood decomposition. Theory predicts that this relationship might be due to the mixture of selection (sampling), complementarity (resource partitioning), and facilitation (physiological activation) effects, whereas the extent to which of these mechanisms working depends on fungal species and combinations. First, in competition between *At* and *Fk*, *Fk* completely replaced *At* in all the experiments, and the weight loss of the blocks was the same as that of pure culture of *Fk*, which has significantly greater decay ability than *At*, representing a selection effect. Similarly, *Tv* replaced *At* in all the experiments, and the weight loss of the blocks showed values close to pure culture of *Tv*. However, in competition between *Og* and *Tv* and between *Fk* and *Tv*, the original fungal distribution seldom changed during incubation, and the weight loss of the blocks showed nearly average values of the pure cultures of the competitors, indicating that none of the above mentioned mechanisms worked in these combinations.

Moreover, in competitions between *At* and *Og* and between *Fk* and *Og*, the weight loss of the blocks showed significantly greater values in some experiments compared with pure cultures of the competitors, which indicates more than just a selection effect. In competition between *At* and *Og*, the competition outcome was complete replacement of *At* by *Og*. The weight loss data of wood blocks where *At* was replaced by *Og*, the weight loss was significantly greater than the pure cultures of *At* and *Og*. In this case, we cannot separate the effects of resource partitioning between *At* and *Og* and the physiological activation of these fungi during competition. Although the weight loss of the blocks in *At* pure cultures was negligible (rather the negative values indicating net increase of weight, which will be discussed later), the species in Xylariaceae caused white rot-like decay; thus, lignin was removed (Abe 1991). The removal of lignin by xylariaceous fungi can increase lignocellulose decay efficiency by successor basidiomycetes (Osono 2003), which leads to resource partitioning between xylariaceous fungi and basidiomycetes. However, the decay activities of *At* and *Og* could be activated within wood blocks they occupied during competition. In GLMM focusing on the wood blocks of *Og* without replacement and zone lines, the number of surface contact with other species’ block showed a significant effect promoting wood decay by *Og*. This effect is a physiological activation of fungi by interaction with other species, not a resource partitioning. Fukasawa et al. (2020) also reported that the wood decay activity of a wood decay basidiomycete was activated by contact with blocks of another basidiomycete species. *At* could also be activated before being replaced by *Og*, but we cannot reveal that because all *At* wood blocks were replaced.

In competition between *Fk* and *Og*, such facilitation effects might be more feasible.

Although *Og* was replaced by *Fk* when the interaction surface was 4 and 8, not a negligible number of *Og* blocks remained after the incubation period when the interaction surface was 24. Similarly, in the four-species competition, the presence of a weaker competitor (*At*) as a neighbor changed the offense target for *Fk* from *Og* to *At* and let *Og* blocks survive. Such intransitivity in competition across wood decay fungi has been reported (Hiscox et al. 2017; O’Leary et al. 2018, 2021) and thought to be an important feature in fungal diversity–decomposition debate (Maynard et al. 2017b). In the present study, *Fk* was the strongest competitor on wood, and its competitive ability increased in the four-species system than in the two-species system (Table S3). The activation of *Fk* in the four-species system was also supported by the result of GLMM, which showed that species number in the system promoted wood decomposition by *Fk*. However, the mechanism underlying this phenomenon remains unclear, but a combination of resource partitioning with *At* and physiological activation in competition with *Og* could be an additive effect. In four-species experiment, *Fk* always replaced *At* completely and competed with *Og* and *Tv*, and the decay promotion in *At* wood blocks replaced by *Fk* was detected only in the four-species system but not in the two-species system (Fig. 7).

Compared with *Og*, *Tv* accelerated wood decay rate when a larger number of their block surfaces were connected to other *Tv* blocks. Therefore, *Tv* had a greater wood decay ability when their wood volume was large. Some previous studies reported that wood decay fungi can accelerate decomposition when their inhabiting wood was small, probably because of the excessive protection cost for a relatively larger surface area for unit volume in smaller wood (Hughes and Boddy 1996; Fukasawa et al. 2020).

Nevertheless, O’Leary et al. (2021) reported that *Tv* showed greater wood decay ability in larger volume of wood. Such discrepancy might also be dependent on the energy allocation strategy of different fungal species. Wood decay fungi can accelerate wood decomposition at the center of their territory when they occupy a large volume of wood possibly to compensate the energy loss at the competition front (Fukasawa et al. 2020).

Lignin and carbohydrate analyses of *Og* and *Fk* wood blocks revealed that L/W decreased when they shared interaction surfaces with *Tv* in two-species experiments. In *Og* blocks, the larger the number of interaction surface, the smaller L/W was recorded. These reductions in L/W might be attributed to two different mechanisms. First, *Og* and *Fk* may shift to more carbohydrate decomposition in competition with *Tv*. Although not significant, interaction with *Tv* and the number of interaction surfaces reduced C/W of *Fk* interacted with *Tv*. The larger amount and mass variation in carbohydrate compared to lignin may musk the small difference in C/W. Second, fungi produce of recalcitrant materials, such as melanin. In *Og* and *Fk*, competition outcomes with *Tv* on wood blocks were deadlock in the two-species experiments. Deadlock interaction may induce zone line formation and melanin production by both competitors (Boddy 2000; Morris et al. 2021). Fungal melanins are recalcitrant and they could be detected as Klason lignin fraction in sulfuric acid digestion, and thus may reduce L/W (Fukasawa et al. 2009). C/W decreased in *Og* blocks competed with *At* in a large number of interaction surfaces. Considering that *Og* accelerated wood decomposition in former *At* blocks that they replaced, the excess energy obtained from this reaction could be reused to accelerate lignin decomposition in the original blocks of *Og*. Although not significant, but *Og* blocks with four interaction surfaces with *At* showed larger L/W than the blocks with smaller interaction surfaces. These results support the findings of O’Leary et al. (2019), who reported that the difference in competition outcomes can affect enzyme production of the competitors: deadlock induced ß-glucosidase activity (i.e., carbohydrate decomposition) but not the manganese peroxidase (MnP) activity (i.e., lignin decomposition). Conversely, replacement induced MnP activity.

## Conclusions

The present study showed clear evidence of the positive effects of fungal species richness and interspecific interactions on wood decomposition. Depending on fungal species combinations, evidence of selection, complementarity, and facilitation effects were also shown. Conversely, clear evidence of decay reduction caused by trade-off allocation to competition cost was not observed. Fungal interspecific competitions are energetically expensive as shown in previous studies (Maynard et al. 2017a,b; Hiscox and Boddy 2017; Hiscox et al. 2010, 2015; O’Leary et al. 2019, 2021). However, whether such competition reduce or accelerate wood decomposition is a different story. The negative correlations between fungal diversity and wood decomposition (mass loss or CO_2_ efflux) observed in the field (Yang et al. 2016; Skelton et al. 2019; Smith and Peay 2021) do not necessarily mean that diverse fungal community reduces wood decomposition. Rather, opposite causality, that is, diverse fungi could co-exist in wood with slow decay rate, is equally possible. Same is true for laboratory incubation experiments evaluating fungal diversity by species richness that remained after incubation (Fukami et al. 2010; Fukasawa and Matsukura 2021). The results of the present study indicated that wood decay fungi accelerate decomposition under competition to compensate the energy cost for competition. Nevertheless, in the cases where deadlock interactions among basidiomycetes are dominant, a greater interaction surface may induce the accumulation of recalcitrant fungal secondary metabolites and may retard wood decomposition in a longer time scale. This relationship is similar to the effect of fungal species richness on decomposition because interaction frequency usually increases with species richness in the field, where large deadwoods are hosting a substantially greater number of fungal species than the present study. Furthermore, the accumulation of fungal recalcitrant materials is important for carbon sequestration in soil organic matter (Nielsen et al. 2011; Liu et al. 2024). We would like to name the negative effect of accumulated recalcitrant fungal metabolites on wood decomposition in BEF context as “accumulated inhibitor hypothesis” and would call for more experimental evaluations.

## Acknowledgements

We are grateful to Yuto Hatanaka, Kosuke Hamano, Xintong Li, Minori Mori, and Tsuyoshi Yamada for their supports in laboratory works. Thanks are extended to Emi Kameoka, Yoshihisa Suyama, Daiki Takahashi, and Naoko Ishikawa for valuable discussions.

## Author contributions

Yu Fukasawa conceived and designed the experiments, received funding, and wrote the original draft. Aoi Chiba performed the experiments, analyzed data, and prepared the figures and tables. Both authors contributed substantially to data interpretation, manuscript writing, revision, and approved the final version.

Conflict of interest None declared.

## Funding

This work was supported by the program of Japan Society for Promotion of Science KAKENHI (No. JP24H02110, JP22H05669) and Kesen Pre-cut Cooperative Association.

## Data availability

The data of this work are deposited in Zenodo (doi:10.5281/zenodo.18676291).

---

**Fig. S1.**
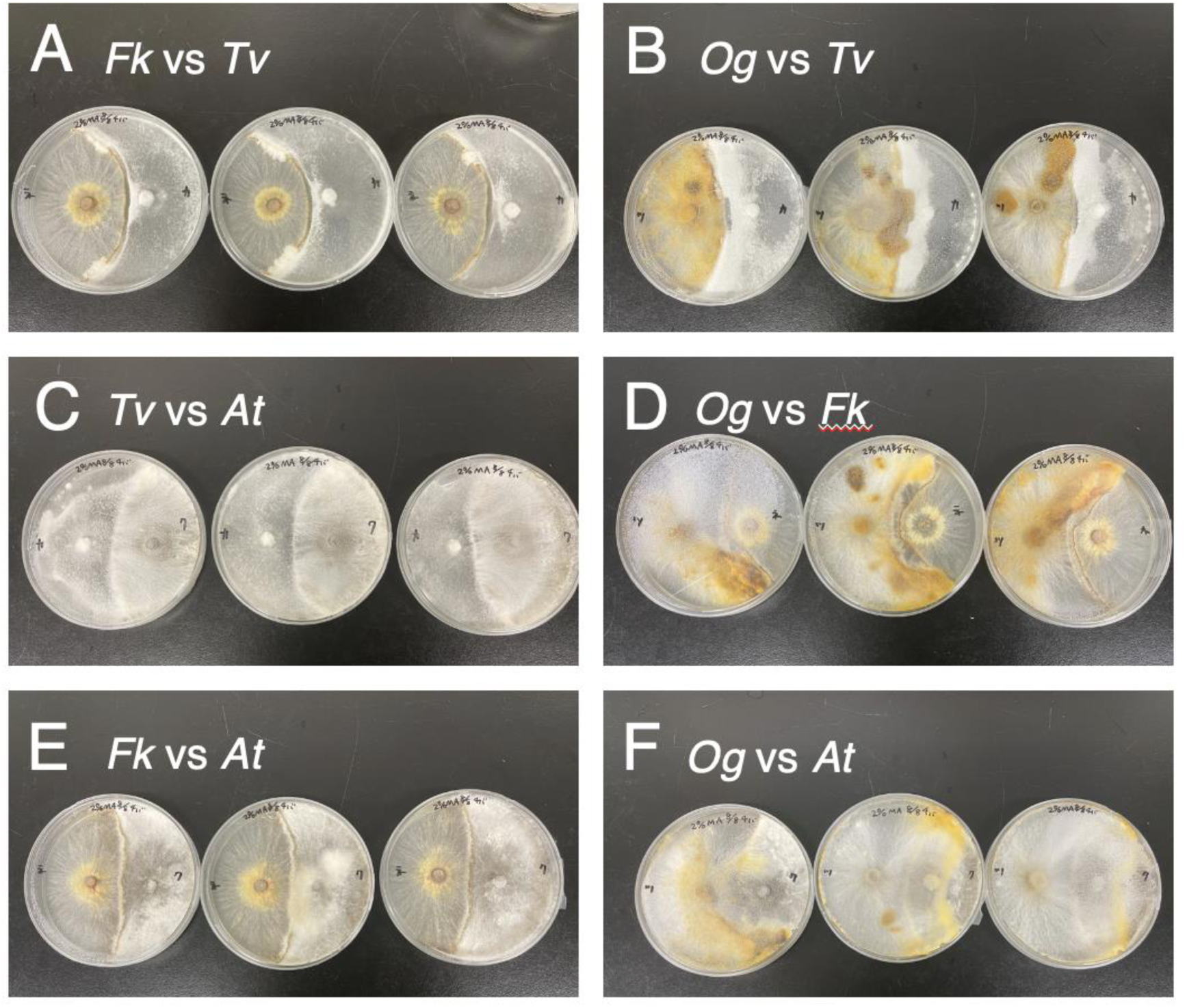
Competition outcome of the four fungal species on 2% Malt extract agar media after incubated at 25°C in the dark for 2–4 weeks. Each photo shows the results of the three replicate dishes. The three replicates showed identical results in all combinations. (A) *Fk* (left) versus *Tv* (right), deadlocked. (B) *Og* (left) versus *Tv* (right), deadlocked. (C) *Tv* (left) replaced *At* (right). (D) *Og* (left) versus *Fk* (right), deadlocked. (E) *Fk* (left) versus *At* (right), deadlocked. (F) *Og* (left) replaced *At* (right).

**Fig. S2.**
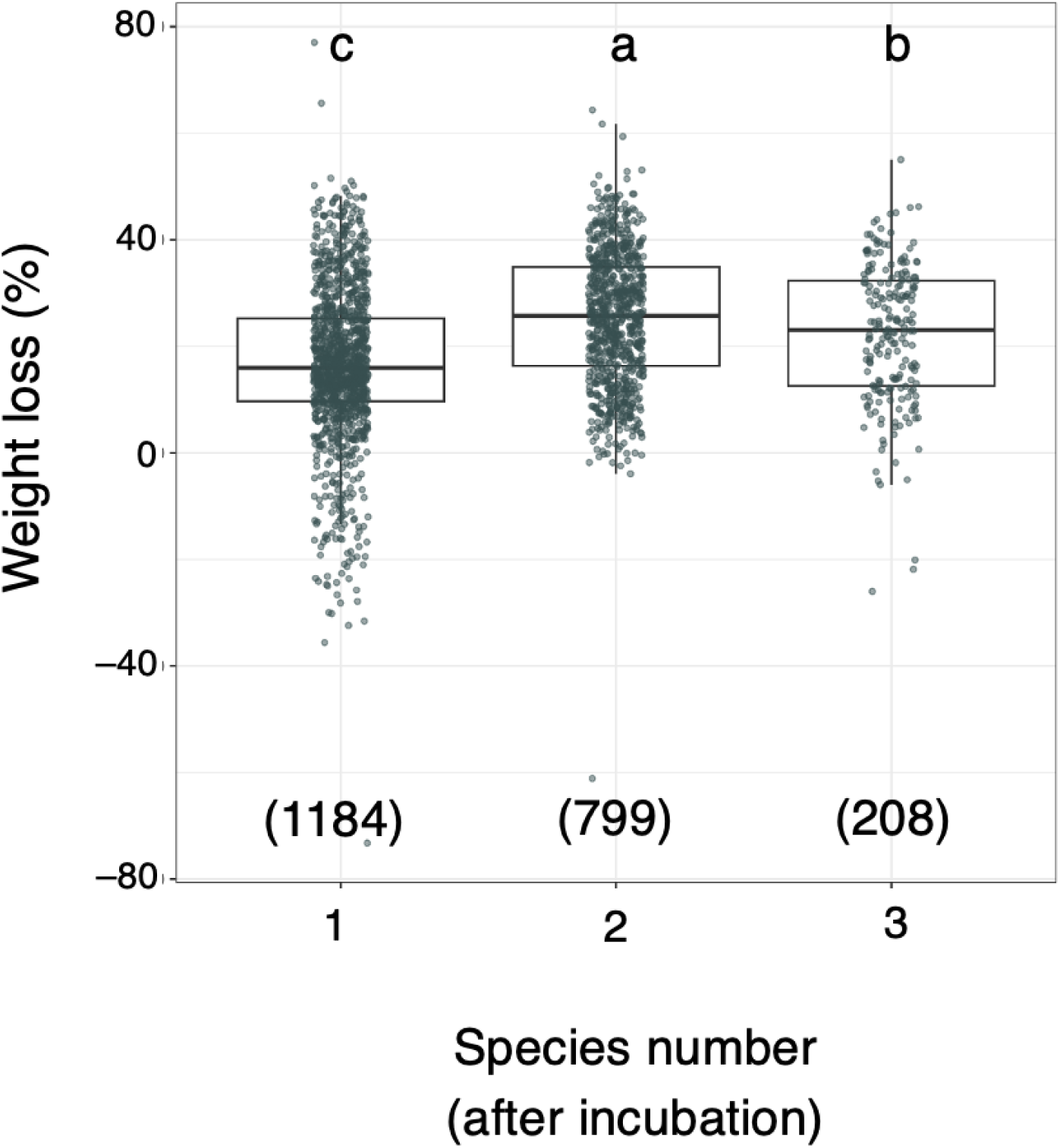
Relationships between weight loss (%) and species number after incubation. The different uppercase and lowercase letters indicate significant difference across the three levels of species number (Steel–Dwass test, *p* < 0.05). Note that none of the four-species experiments retained original four species after the incubation. The numbers enclosed in parentheses are the sample number.

**Fig. S3.**
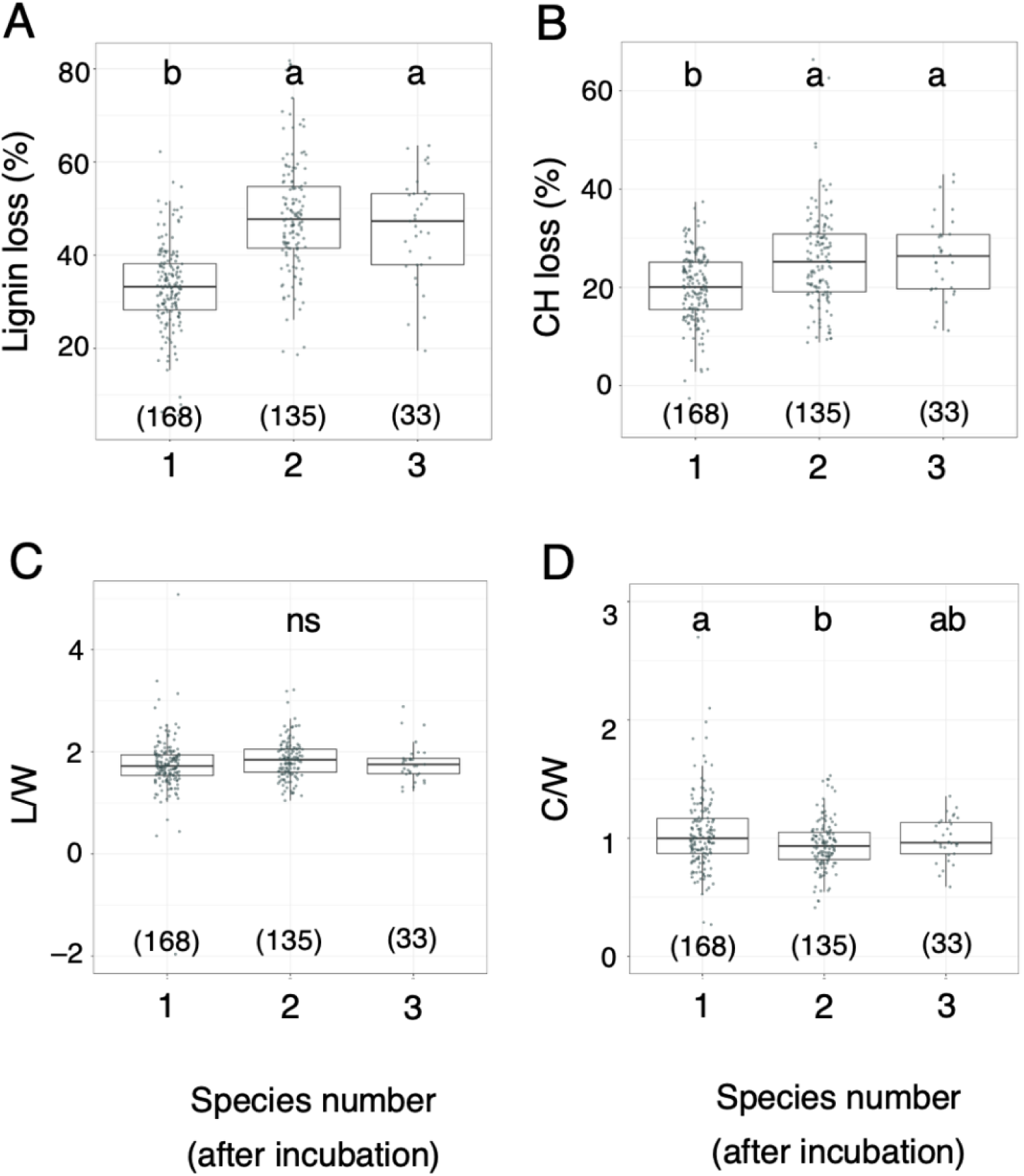
Effects of fungal species number after the incubation on *Fk* blocks’ weight loss (%) of lignin (A) and carbohydrate (B), and relative weight loss of lignin (C) and carbohydrate (D) against weight loss of wood. Data of 336 *Fk* blocks without replacement and zone lines were used. Two outliers in each L/W and C/W were out of the plots. The different lowercase letters indicate significant difference across the blocks with different fungal species number (Steel–Dwass test, *p* < 0.05). ns, no significant difference. Note that none of the four-species experiments retained original four species after the incubation. The numbers enclosed in parentheses are the sample number.

**Fig. S4.**
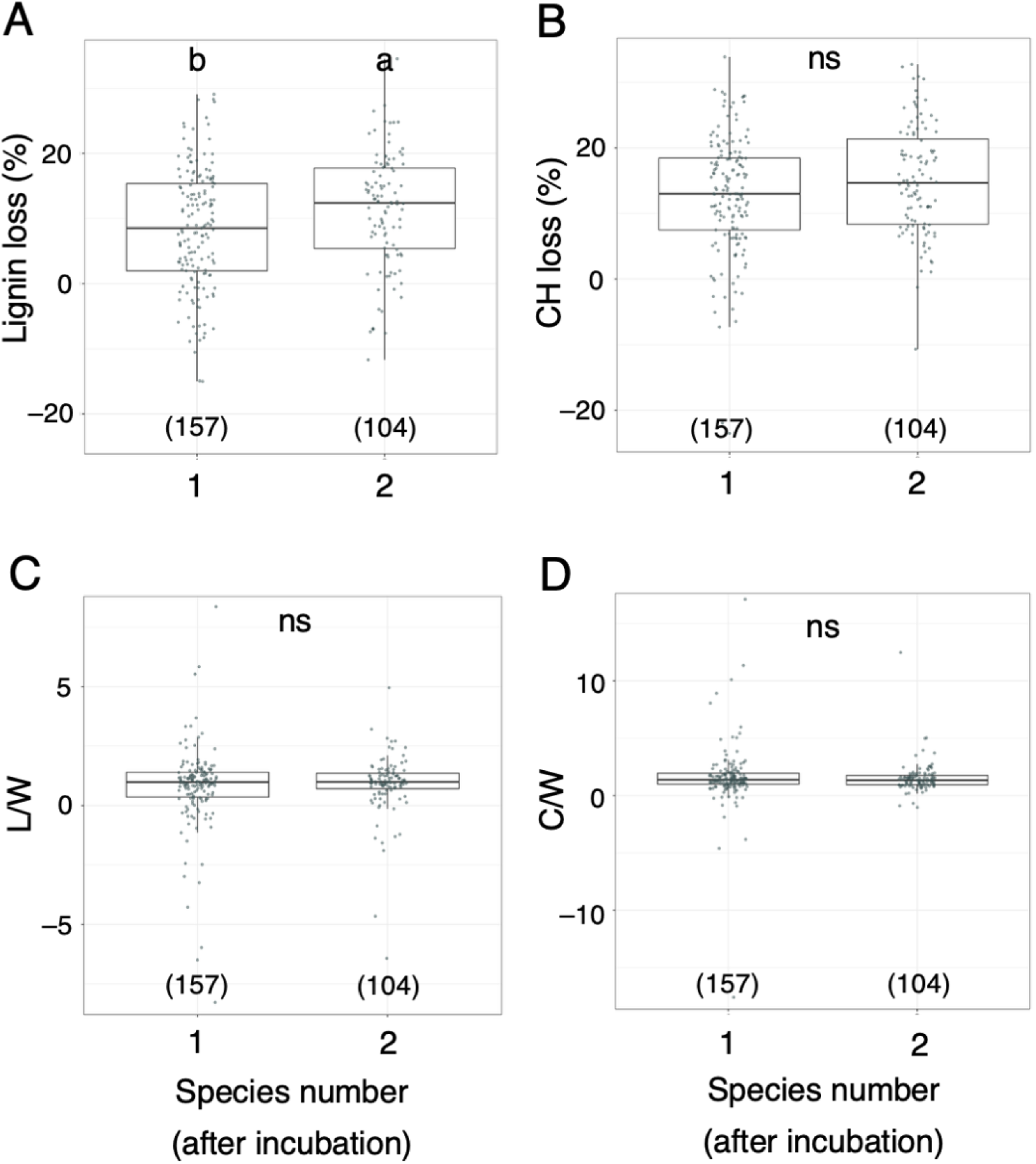
Effects of fungal species number after the incubation on *Og* blocks’ weight loss (%) of lignin (A) and carbohydrate (B), and relative weight loss of lignin (C) and carbohydrate (D) against weight loss of wood. Data of 261 *Og* blocks without replacement and zone lines were used. Four outliers in each L/W and C/W were out of the plots. The different lowercase letters indicate significant difference across the blocks with different fungal species number (Steel–Dwass test, *p* < 0.05). ns, no significant difference. The numbers enclosed in parentheses are the sample number.

**Table S1.** Fungal strains used in this study.

| Specues | Code | NBRC# | Phylla | Wood inhabiting strategy | Decay type | Reference |
| --- | --- | --- | --- | --- | --- | --- |
| *Annulohypoxylon truncatum* | *At* | 115313 | Ascomycota | Endophyte/primary colonizer | Soft/White | Abe 1991; Helaly et al. 2018 |
| *Fuscoporia koreana* | *Fk* | 115317 | Basidiomycota | Secondary colonizer | White | Banik et al. 2024 |
| *Omphalotus guepiniiformis* | *Og* | 115315 | Basidiomycota | Secondary colonizer | White | Fukasawa et al. 2010 |
| *Trametes versicolor* | *Tv* | 115316 | Basidiomycota | Early secondary colonizer | White | Hiscox et al. 2010 |

**Table S2.** Competition outcomes of the four fungal species on 2% malt extract agar media.

| Species 1 | Species 2 |  |  |
| --- | --- | --- | --- |
|  | *Tv* | *Fk* | *At* |
| *Og* | D (3) | D (3) | R (3) |
| *Tv* | – | D (3) | R (3) |
| *Fk* | – | – | D (3) |

D, species 1 and 2 deadlocked. R, species 1 replaced the species 2.
The numbers in parenthesis shows the number of replicate dishes showing the same results.

**Table S3.** Number of the wood blocks from which the fungal species were reisolated after the incubation experiment.

| Experiment |  | Wood blocks |  |  |  |  |  |
| --- | --- | --- | --- | --- | --- | --- | --- |
| Species number | Interaction surface | *Tv* | *Fk* | *Og* | *At* | Fail † | Total |
| 2 | 4 | 160 | 199 | 120▲ | 0 | 1 | 480 |
|  | 8 | 156 | 200 | 112 | 0 | 12 | 480 |
|  | 24 | 160 | 182▽ | 136▲ | 0 | 2 | 480 |
| 4 | 8 | 98 | 123▲ | 16▽ | 0 | 3 | 240 |
|  | 24 | 114.5▲ | 104 | 19.5▽ | 0 | 2 | 240 |

†No fungi were reisolated from the block or contamination.
If two species were reisolated from one block, each species got 0.5 point.
Results of chi-square test was shown as ▲ if the recorded value was significantly larger than expected value and as ▽ if the recorded value was significantly smaller than the expected value ( $p < 0.05$ ).

**Table S4.** GLMM results illustrating the relationships between dry weight of wood blocks and factors.

| Factor | *Tv* | *Og* | *Fk* |
| --- | --- | --- | --- |
| Number of surface contact with other species' block | – | –0.028*** | – |
| Number of surface contact with same species' block | –0.024* | – | – |
| Fungal species number before experiment § | – |  | –0.097*** |
| Water content | – | –0.268** | –0.172*** |

Data of wood blocks from which original colonized fungus was reisolated were used for the models.
Only estimated coefficients selected in best models were shown.
*A. truncatum* model was not tested because all the blocks were replaced by other species.
§Models used fungal species number after experiment gave similar results (see Table S1).
\*, $p < 0.05$ ; \*\*, $p < 0.01$ ; \*\*\*, $p < 0.001$ .

**Table S5.** GLMM results illustrating the relationships between dry weight of wood blocks and factors.

| Factor | *Tv* | *Og* | *Fk* |
| --- | --- | --- | --- |
| Number of surface contact with other species' block | – | –0.028*** | – |
| Number of surface contact with same species' block | –0.024* | – | – |
| Fungal species number after experiment § | – |  | –0.095*** |
| Water content | – | –0.268** | –0.172*** |

Data of wood blocks from which original colonized fungus was reisolated were used for the models.
Only estimated coefficients selected in best models were shown.
*A. truncatum* model was not tested because all the blocks were replaced by other species.
§Models used fungal species number after experiment gave similar results (see Table S1).
\*, $p < 0.05$ ; \*\*, $p < 0.01$ ; \*\*\*, $p < 0.001$ .

## Notes

https://zenodo.org/records/18676291

### Competing Interest Statement

The authors have declared no competing interest.

